# Both density- and frequency-dependent effects determine plant growth in a dune heath ecosystem

**DOI:** 10.1101/2024.09.12.612621

**Authors:** Christian Damgaard, Mathias Emil Kaae, Jacob Weiner, Jesper Leth Bak

**Author notes:** Correspondence: Christian Damgaard.

## Abstract

We tested the hypothesis that both density- and frequency-dependent interactions play important roles in determining plant growth in a dune heath ecosystem at several levels of available nitrogen. Plant growth was measured using the pin-point method in a five-block experiment with four nitrogen levels. To maximize statistical power, we used only three taxonomic groups: *Calluna vulgaris*, *Avenella flexuosa* (the two most dominant species), and all other vascular plant species together. The results show that both Lotka-Volterra type interspecific competition and frequency-dependency play significant roles in determining the growth of the species in the community. Significant interspecific density-dependent competition was observed in 4 out of the 6 possible cases. Nitrogen addition increased the competitive effect of *C. vulgaris* on the growth of the other species. Both *C. vulgaris* and *A. flexuosa* showed frequency-dependent positive feedback dynamics on growth when they were relatively dominant at the plot scale, and this effect increased with added nitrogen. In plots with added nitrogen, there was a beneficial effect of being relatively rare on the growth of the group of other species. The study highlights the importance of the combined effects of density and frequency dependency in determining plant growth.

## Introduction

To develop predictive models of ecosystems in changing environments, it is important to understand the mechanisms that govern plant community dynamics and quantify their relative importance. Historically, the most important mechanisms for understanding plant community dynamics are competition for resources, e.g. light, water and soil nutrients that are necessary for plant growth (Goldberg et al. 1990). Species coexistence has primarily been examined with density-dependent Lotka-Volterra type models, where population growth is affected by intra- and interspecific population densities, and investigating the conditions for invasion when a species is rare (Adler et al. 2018; Barabás et al. 2018; Chesson 1994; 2000). This body of theory has been referred to as “modern coexistence theory”. But plant communities may also be regulated by frequency-dependent mechanisms, where plant growth is regulated by the frequency of the species within the community. There has been increasing awareness that frequency-dependent mechanisms may play important roles in the dynamics of plant communities (Bever et al. 1997; Chisholm and Fung 2020; Connell et al. 1984; Suding et al. 2024). Such frequency-dependency may be caused by numerous mechanisms, such as species-specific soil-plant interactions that hinder local establishment and growth of conspecific dominant plant species, or because relatively low population sizes of natural enemies of rare species may give them an advantage (Heinen et al. 2020; Mazzoleni et al. 2015a; Mazzoleni et al. 2015b; van der Putten et al. 2013), or dominant plant species may be favored by positive feedback mechanisms, e.g. by resource monopolizing strategies (van der Putten et al. 2013). It is surprising that many studies demonstrate the importance of frequency dependence for coexistence, but it is completely ignored in “modern coexistence theory”. It has also been argued that higher-order interactions among species are important (Singh and Baruah 2020), and that intraspecific genetic variation may play a role in the outcome of species interactions (Ehlers et al. 2016). Generally, the relative importance of different mechanisms in regulating plant community dynamics in different ecosystems is still an open question (e.g. De Long et al. 2023).

Dune heathland ecosystems are natural or semi-natural habitats situated on leached, nutrient-poor, sandy soils. The vegetation is adapted to withstand the harsh, windy environment and is dominated by dwarf shrubs, e.g. *Calluna vulgaris* (heather), *Empetrum nigrum* (crowberry), *Vaccinium uliginosum* (bog bilberry), and graminoids, especially *Avenella flexuosa* (wavy hair-grass), *Carex arenaria* (sand sedge), *Corynephorus canescens* (grey hair-grass), and *Festuca ovina (*sheep fescue). Bryophytes and lichens are also common. Dwarf shrubs and grasses often coexist in a dynamic patchy spatial configuration. The cyclical successional processes leading to such a patchy distribution of dwarf shrubs and grasses was described by Watt (1947), a key feature of which is the life cycle of *Calluna vulgaris* plants, which senescence when they are 30–40 years old (Gimingham 1988; Watt 1955).

One of the essential abiotic factors that regulate plant growth and community dynamics is the level of available nitrogen (Craine 2009). Nitrogen deposition from the combustion of fossil fuels and animal husbandry is known to play an important role in current heathland ecosystems (Alonso et al. 2001; Bähring et al. 2017; Damgaard 2019a; 2019b; Jones and Power 2012; Southon et al. 2013). If nitrogen input is sufficiently high, especially after the vegetation is opened by burning or beetle infestation, grass species may outcompete ericoid dwarf shrubs and the dwarf shrub vegetation may be replaced by acid grassland vegetation or, in the absence of management, encroachment by trees (Alonso et al. 2001; Heil and Bobbink 1993; van Voorn et al. 2016). Under increased nitrogen availability, *A. flexuosa* outcompetes *C. vulgaris*, which is adapted to nitrogen-limited environments in which its slowly decaying litter is decomposed by ericoid mycorrhiza and the nitrogen in its litter is absorbed by its roots (Diemont and Heil 1984; Johansson 2001; Smith and Read 2008; Talbot et al. 2008). The positive feedback between heather and ericoid mycorrhiza may lead to frequency-dependency in nitrogen limited environments, where the competitive growth of *C. vulgaris* is accelerated when it is dominant (Aerts et al. 1990).

The role of competition in determining plant growth and therefore community dynamics is investigated empirically by modelling the observed net negative plant-plant interactions (“competition in the broad sense” sensu Weiner 1993) using Lotka-Volterra-type competition models (e.g Adler et al. 2018; Damgaard 1998; 2005; Law and Dieckmann 2000). Although the use of net effects in the modelling of species interaction has been criticized, it is still the best currently available tool we have for describing inter- and intra-specific interactions in communities. Hypothesizing that frequency dependence plays an important role in plant community dynamics, Damgaard and Weiner (2021) added frequency dependent competition to a Lotka-Volterra-type competition model. Their model was fitted to empirical growth data using a hierarchical Bayesian framework, in which the possible role of unmeasured variables that influence individual plant performance and measurement errors was accounted for (Fig. 1). The inclusion of frequency dependence improved the model fit significantly.

**Fig. 1.**
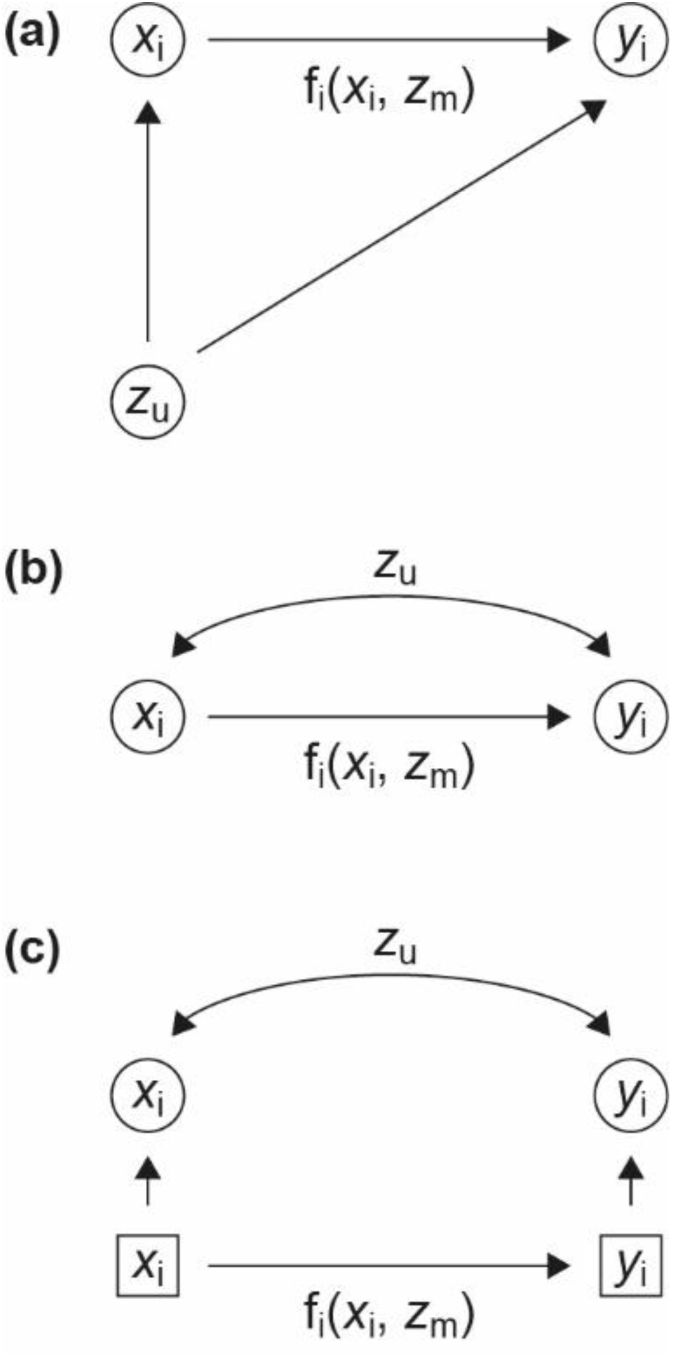
Flowcharts illustrating components of the model. (a) The effect of an unmeasured variable (*Z*_*u*_) on individual plant performance at both an early (*x*_*i*_) and later growth stage (*y*_*i*_) of plant species *i*. The competitive growth process is modelled by the function, *f*_*i*_, which depends on *x*_*i*_ and possibly on some measured environmental variables (*Z*_*m*_). (b) The effect of the unmeasured variable affecting both earlier and later growth stages is modelled as the part of the covariance between the early and later growth stages that is not explained by the independent factors in the competition model. (c) Hierarchical model in which true but unknown factors affecting plant performance are modelled by latent variables (squares) and measurement errors are separated from process errors. Data are denoted by circles.

Here, we test the hypotheses that (1) both density- and frequency-dependent competition play important roles in determining plant growth in a dune heath ecosystem, and (2) that the relative importance of these different mechanisms depends on the amount of available nitrogen. Growth models in which the effect of density- and frequency-dependent competition are parameterized and fit to pin-point cover data sampled in both early and late summer over a three-year period. The competitive growth of *C. vulgaris* and *A. flexuosa*, and all other higher plants grouped together are then modeled as a function of the cover in early summer (Damgaard and Weiner 2021).

## Material and Methods

### Competitive growth data

At a dune heath site, Vust Hede (Fig. 2), with relatively low background nitrogen deposition (9 kg N ha^-1^ year^-1^), five experimental blocks were established in 2018 (Kaae et al. 2024). Each block had four treatment plots with added nitrogen (0, 5, 10, 25 kg N ha^-1^ year^-1^). At the sites, there were natural populations of roe deer, red deer and fallow deer in an estimated combined density of 9.95 ha^-1^ day^-1^ (Appendix A), and herbivory by the deer population was shown to affect plant species composition (Kaae et al. 2024). In the summer of 2020, approximately 70-80% of the heather plants were infested by the heather beetle (*Lochmeae suturalis*), leading to discoloring of leaves and reduction in growth (Kaae et al. 2024).

**Fig. 2.**
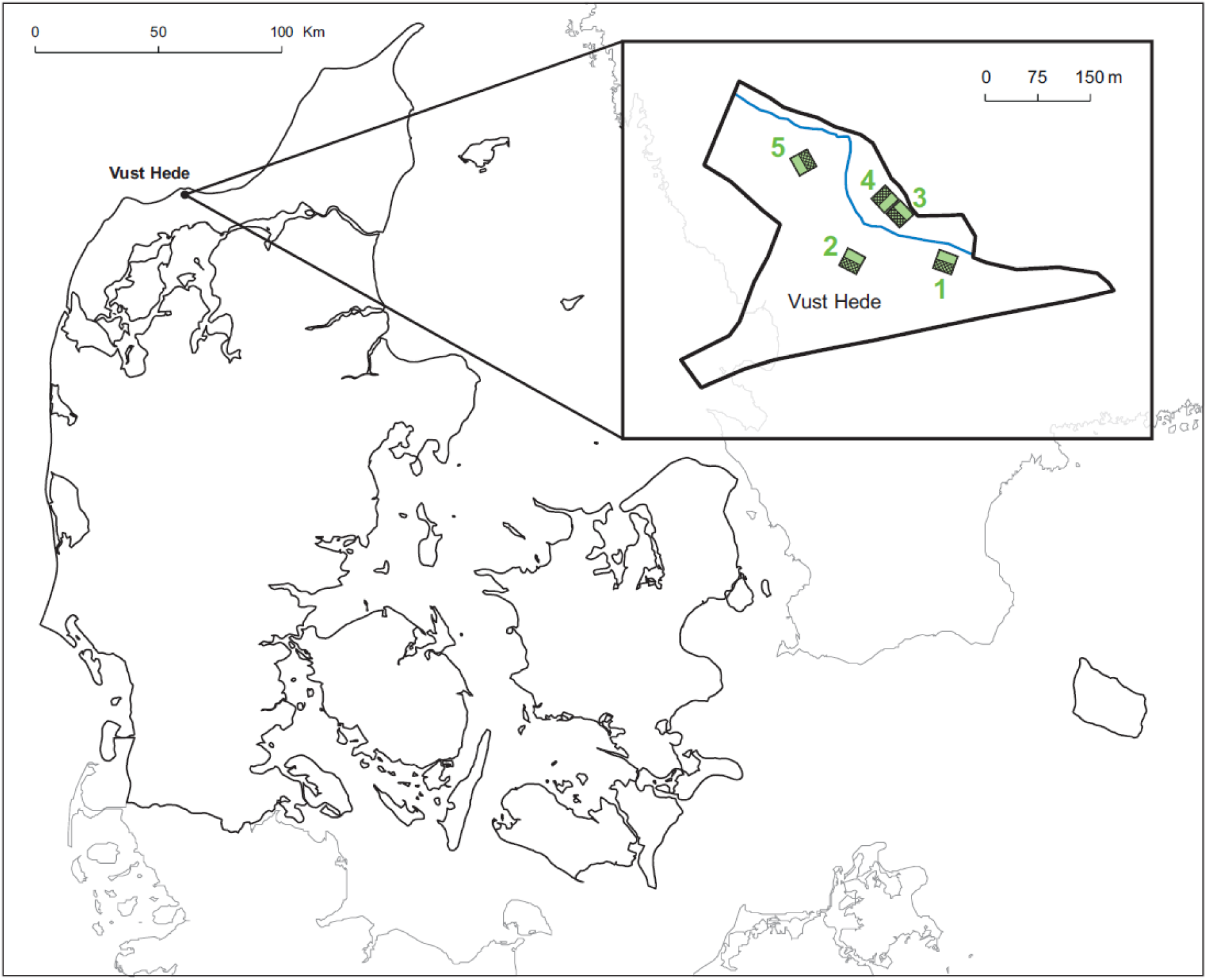
Map showing the location of Vust Hede (57°7′23.412″ N, 9°0′42.443″ E). The rectangles are the blocks (1-5) with experimental manipulations (Kaae et al. 2024).

To study the species interactions on plant growth, a 0.5 m x 0.5 m quadrat was permanently outlined within each plot. In the quadrat, plant abundance was measured non-destructively by the “pin-point” (also called “point intercept”) method (Kent and Coker 1992) using a pin-point frame with the same dimension as the quadrat and with 25 pin-positions regularly placed at distances of 10 cm. At each position, a pin with a diameter of 0.5 mm was passed vertically through the vegetation, and the number of contacts to each species was recorded. The sampling was performed in early (June) and late summer (August to October) in the three years 2019, 2020, and 2021 (Kaae et al. 2024).

The pin-point method provides estimates of two important plant ecological variables: plant cover and vertical density, which are well suited for studying species interactions in natural and semi-natural herbaceous perennial plant communities, where it is difficult to distinguish individual genets (Damgaard 2011; Damgaard et al. 2009). The cover of a specific plant species is defined as the proportion of pins in the grid that touch the species, thus, plant cover measures the cover of the plant species when it is projected onto the two-dimensional ground surface. The vertical density is defined as the number of times a pin touches a specific species, and this measure is correlated with plant biomass (Jonasson 1983; 1988; Kaae et al. 2024).

In the subsequent analysis, the observed plant taxa were grouped into *C. vulgaris*, *A. flexuosa* grass and an aggregated group of all other vascular plant species. The pinpoint data is available in an electronic supplement (Appendix B)

### Model

The effect of species interactions on plant growth was analyzed in a hierarchical model (Fig. 1c) by modelling how the vertical density of *C. vulgaris*, *A. flexuosa* and the other species at the end of the growing season depended on the cover at the beginning of the growing season. The underlying assumption is that the measure of vertical density at the end of the growing season as a function of the cover at the beginning of the growing season may be used as a measure of population growth (Damgaard 2011; Damgaard et al. 2009).

We used a modified version of the model in Damgaard and Weiner (2021),

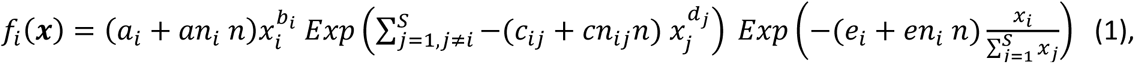

where *f*_*i*_(*x*) is the expected vertical density of species *i* at the end of the growing season, and *x*_*i*_ is the cover of species *i* at the beginning of the growing season. The first term models the intraspecific density dependence (i.e. if the species were in a monospecific population).

The second term models the effect of interspecific density dependence (as in the Lotka-Volterra models), and the third term models the effect of frequency-dependency within the plant community. Thus, the density-dependent and frequency-dependent effects are partitioned and estimated in separate terms. More specifically, the parameters *a*_*i*_, *an*_*i*_ and *b*_*i*_ model the expected vertical density as a function of cover and nitrogen (*n*) in the absence of inter-specific density-dependent and frequency-dependent Effects, while *c*_*ij*_, *cn*_*ij*_ and *d*_*j*_ model the inter-specific interactions of species *j* on the vertical density of species *i*. If (*c*_*ij*_ + *cn*_*ij*_ *n*) > 0, there is evidence of competitive inter-specific interactions, otherwise there is evidence of facilitative inter-specific interactions. The parameters *e*_*i*_ and *en*_*i*_ model the frequency-dependent effect of species *i*. If (*e*_*i*_ + *en*_*i*_ *n*) = 0, then there is no evidence of frequency-dependence, and the population growth model (1) simplifies to a Lotka-Volterra competition model. If (*e*_*i*_ + *en*_*i*_ *n*) > 0, then rare species have an advantage over dominant species and, conversely, if (*e*_*i*_ + *en*_*i*_ *n*) < 0, then dominant species have an advantage over rare species. The parameters *b*_*i*_ and *d*_*j*_ model possible additional non-linearity in the effect of cover on vertical density with a domain in the interval (0.5, 2).

The measurement errors of cover and vertical density have previously been assumed to be binomially distributed and generalized-Poisson distributed, respectively (Damgaard et al. 2014). However, since we wanted to model the covariance between the early and later growth stages to account for the possible effect of unmeasured variables (Rinella et al. 2020), both distributions are approximated by standard normal distributions, where (i) the observed pin-point cover measure, *y*, in a pin-point frame with *n* pin-positions and an expected cover *q* is transformed to 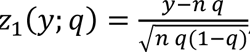, and (ii) the observed pin-point vertical density measure, *vd*, with an expected value λ and with a species-specific scale parameter υ is transformed to 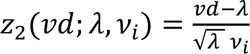. The measurement error of the standardized cover and vertical density measures is consequently modelled as

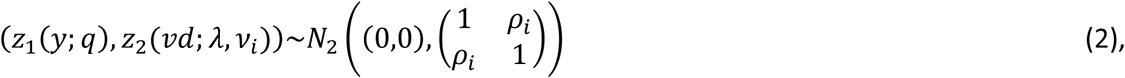

where ρ_*i*_ is a species-specific correlation coefficient in the domain (−1, 1; (Damgaard and Weiner 2021). If ρ_*i*_ = 0, then the measurement errors of the standardized cover and vertical density measures are assumed to be uncorrelated, i.e. the competition model, not additional unmeasured variables, accounts for the covariance between the early and later growth stages.

The structural uncertainty of the competitive growth model was assumed to be normally distributed and modelled as *vd*_*i*_∼*N*(*f*_*i*_(*x*), σ_*i*_), where σ_*i*_ is the species-specific process error (see Table 1).

**Table 1.**
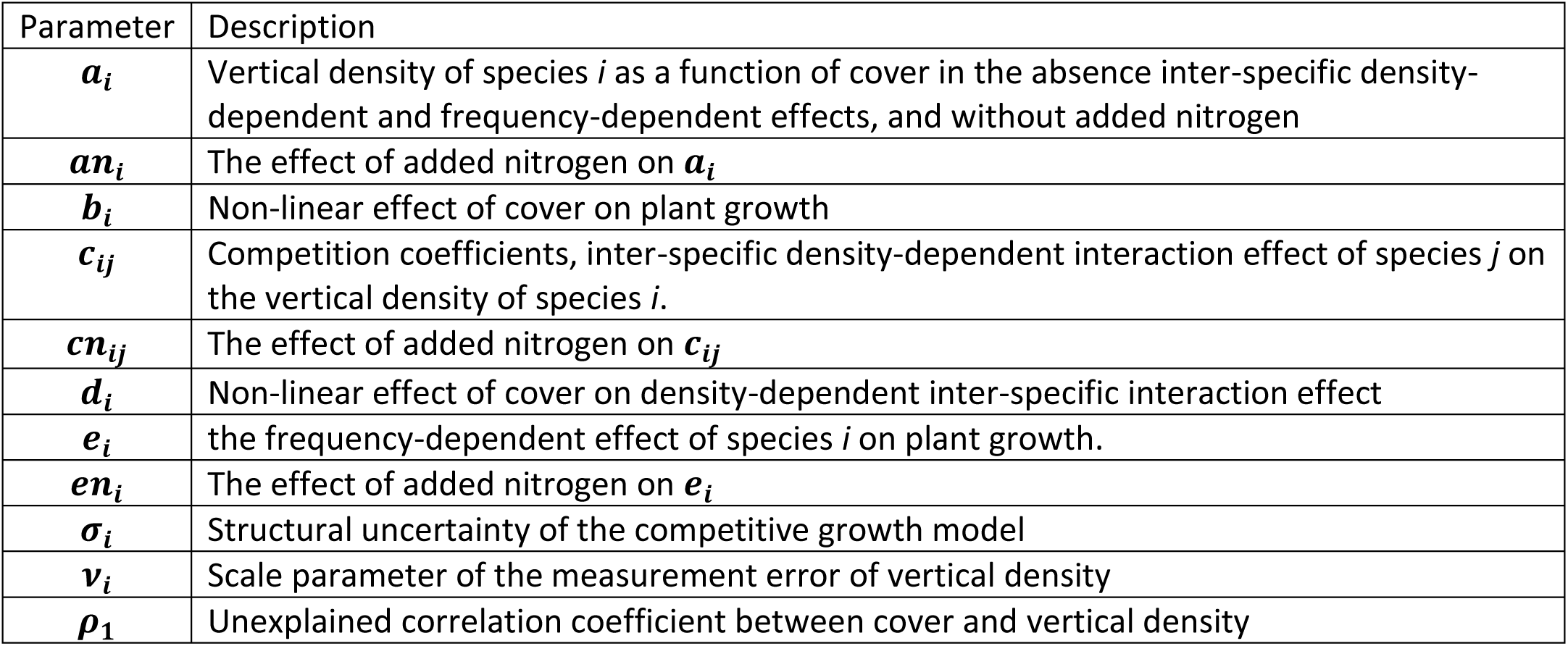
Parameter explanations.

| Parameter | Description |
| --- | --- |
| $a_i$ | Vertical density of species $i$ as a function of cover in the absence inter-specific density-dependent and frequency-dependent effects, and without added nitrogen |
| $an_i$ | The effect of added nitrogen on $a_i$ |
| $b_i$ | Non-linear effect of cover on plant growth |
| $c_{ij}$ | Competition coefficients, inter-specific density-dependent interaction effect of species $j$ on the vertical density of species $i$. |
| $cn_{ij}$ | The effect of added nitrogen on $c_{ij}$ |
| $d_i$ | Non-linear effect of cover on density-dependent inter-specific interaction effect |
| $e_i$ | the frequency-dependent effect of species $i$ on plant growth. |
| $en_i$ | The effect of added nitrogen on $e_i$ |
| $\sigma_i$ | Structural uncertainty of the competitive growth model |
| $v_i$ | Scale parameter of the measurement error of vertical density |
| $\rho_1$ | Unexplained correlation coefficient between cover and vertical density |

### Estimation

We treat the three years of competitive growth data from each of the plots as independent events because i) the within-plots among-year covariation is expected to be minimal, since we model the plant growth of each year, and ii) considering plot as a random effect is not compatible with modelling covariance between early and late plant growth.

The joint Bayesian posterior probability distribution of the parameters in the model was calculated using Markov Chain Monte Carlo (MCMC; Metropolis-Hastings) simulations of 900,000 iterations with a burn-in period of 800,000 iterations and normal candidate distributions (Carlin and Louis 1996). The prior probability distributions of all parameters and latent variables were assumed to be uniformly distributed within their specified domains under the additional constraints that 0 < *a*_*i*_ < 500,−5 < *c*_*ij*_ < 5, −5 < *e*_*i*_ < 5, 0.5 < ρ_*i*_ < 8, and −0.5 < ρ_*i*_ < 0.8, except for σ_*i*_, which was assumed to be inverse gamma distributed with parameters (0.1, 0.1).

Plots of the deviance and trace plots of parameters were inspected to check the fitting and mixing properties of the sampling procedure. Residual plots of the vertical density for each species were inspected to check the fitting properties of the different models. The statistical inferences were assessed using the calculated credible intervals, i.e. 95% percentiles of the marginal posterior distribution of the parameters and the probabilities that the marginal posterior distributions of the parameters were either above or below zero. All calculations were done using Mathematica (Wolfram 2019). The software code and all results are given in the electronic supplement (Appendix B).

### Prediction of relative vertical density

To investigate the estimated effects of both density- and frequency-dependent competition on plant growth (vertical density) at different cover values of the three species and nitrogen levels, the mean relative predicted vertical density was calculated by inserting samples from the joint posterior distribution of the model parameters into equation 1 and dividing by the cover of the species, *f*_*i*_(*x*)/*x*_*i*_.

## Results

The MCMC iterations of the plant growth model converged with acceptable mixing properties, although there was sizeable covariation among parameters (Appendix C). Based on the residual plots (Fig. S1), the fit of the model was judged to be acceptable. A summary of the marginal posterior probability distributions of all parameters is shown in Table 2.

**Table 2.** Percentiles of the marginal posterior probability distribution of the model parameters. The subscripts indicate the taxa: 1 - *Calluna vulgaris*, 2 - *Avenella flexuosa*, and 3 - all other vascular plant species. The probability that the marginal distribution is above zero is given in the last column, where significant values are indicated in bold.

| Parameter | 2.5% | 50% | 97.5% | P(X > 0) |
| --- | --- | --- | --- | --- |
| $a_1$ | 60.354 | 66.953 | 72.629 | - |
| $a_2$ | 12.062 | 16.665 | 23.747 | - |
| $a_3$ | 37.188 | 62.952 | 68.653 | - |
| $an_1$ | -1.248 | -0.588 | 0.351 | 0.208 |
| $an_2$ | -0.641 | -0.405 | 0.697 | 0.100 |
| $an_3$ | 0.228 | 0.961 | 1.910 | **0.993** |
| $b_1$ | 1.541 | 1.820 | 1.987 | - |
| $b_2$ | 0.939 | 1.459 | 1.941 | - |
| $b_3$ | 1.635 | 1.916 | 1.996 | - |
| $c_{12}$ | 2.086 | 3.127 | 4.295 | **>0.999** |
| $c_{13}$ | 0.817 | 1.385 | 1.817 | **>0.999** |
| $c_{21}$ | -1.036 | 0.114 | 1.691 | 0.541 |
| $c_{23}$ | 0.596 | 1.490 | 2.892 | **>0.999** |
| $c_{31}$ | 0.222 | 0.651 | 1.568 | **>0.999** |
| $c_{32}$ | -0.719 | 0.117 | 0.872 | 0.636 |
| $cn_{12}$ | -0.118 | 0.008 | 0.228 | 0.548 |
| $cn_{13}$ | -0.006 | 0.022 | 0.079 | 0.877 |
| $cn_{21}$ | -0.256 | -0.058 | 0.086 | 0.344 |
| $cn_{23}$ | -0.074 | 0.065 | 0.136 | 0.709 |
| $cn_{31}$ | 0.118 | 0.236 | 0.346 | **>0.999** |
| $cn_{32}$ | -0.067 | 0.005 | 0.178 | 0.532 |
| $d_1$ | 0.826 | 1.511 | 1.962 | - |
| $d_2$ | 1.157 | 1.656 | 1.986 | - |
| $d_3$ | 1.552 | 1.823 | 1.988 | - |
| $e_1$ | -0.260 | -0.202 | -0.133 | **<0.001** |
| $e_2$ | -0.663 | -0.467 | 0.013 | **0.027** |
| $e_3$ | 0.859 | 1.282 | 1.466 | **>0.999** |
| $en_1$ | -0.124 | -0.104 | -0.095 | **<0.001** |
| $en_2$ | -0.315 | -0.250 | -0.176 | **<0.001** |
| $en_3$ | -0.054 | -0.005 | 0.006 | 0.244 |
| $\sigma_1$ | 0.060 | 0.092 | 0.138 | - |
| $\sigma_2$ | 0.036 | 0.080 | 0.145 | - |
| $\sigma_3$ | 0.042 | 0.333 | 1.059 | - |
| $\nu_1$ | 7.437 | 7.889 | 7.996 | - |
| $\nu_2$ | 7.679 | 7.944 | 7.998 | - |
| $\nu_3$ | 7.796 | 7.960 | 7.999 | - |
| $\rho_1$ | -0.313 | -0.048 | 0.283 | 0.372 |
| $\rho_2$ | -0.406 | -0.062 | 0.341 | 0.390 |
| $\rho_3$ | 0.671 | 0.773 | 0.799 | **>0.999** |

The vertical density in the absence of interspecific species interaction was modelled by the parameters *a*_*i*_, *an*_*i*_ and *b*_*i*_. The vertical density was an increasing function of added nitrogen, only for all the group of other vascular plant species (Table 2, *an*_3_).

The competitive effect of *A. flexuosa* and all other vascular species on within-season growth of *C. vulgaris* was relatively large without added nitrogen (Table 2, *c*_12_, *c*_13_), but there was no significant effect of the nitrogen treatment (Table 2, *cn*_12_, *cn*_13_). Similarly, the competitive effect of all other vascular species on *A. flexuosa* was relatively large without added nitrogen (Table 2, *c*_23_), and there was no significant effect of the nitrogen level (Table 2, *cn*_23_). The competitive effect of *C. vulgaris* on all other vascular species was significantly positive without added nitrogen (Table 2, *c*_31_), and this effect increased significantly with the level of added nitrogen (Table 2, *cn*_31_). There was no significant competitive effects of *A. flexuosa* on the growth of all other vascular species (Table 2, *c*_32_, *cn*_32_), or of *C. vulgaris* on the growth of *A. flexuosa* (Table 2, *c*_21_, *cn*_21_).

There were significant frequency-dependent effects. For *C. vulgaris* and *A. flexuosa*, there were significant positive effects of being relatively dominant at the plot scale (Table 2, Fig. 3, *e*_1_, *e*_2_), and this effect increased significantly with added nitrogen (Table 2, Fig. 3, *en*_1_, *en*_2_). On the other hand, all other vascular species were significantly positively affected when relatively rare in the plots (Table 2, *e*_3_, *en*_3_).

**Fig. 3.**
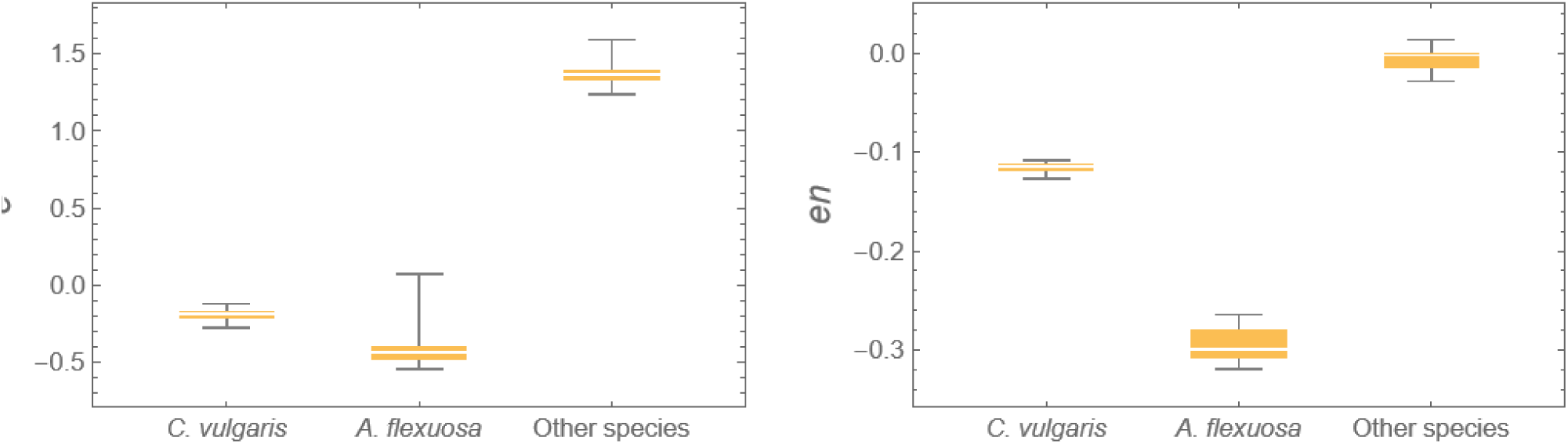
Marginal posterior distributions of the parameters *e*_*i*_ and *en*_*i*_that model frequency-dependent growth. If (*e*_*i*_ + *en*_*i*_ *n*) > 0, then rare species have an advantage over dominant species and, conversely, if (*e*_*i*_ + *en*_*i*_ *n*) < 0, then dominant species have an advantage over rare species.

The mean relative predicted vertical density of both *C. vulgaris* and *A. flexuosa* increased with cover (Fig. 4), indicating that the frequency-dependent positive feedback mechanisms are important for the plant community dynamics in heathland ecosystems. The predicted increases were relatively independent of the amount of added nitrogen (results not shown).

**Fig 4.**
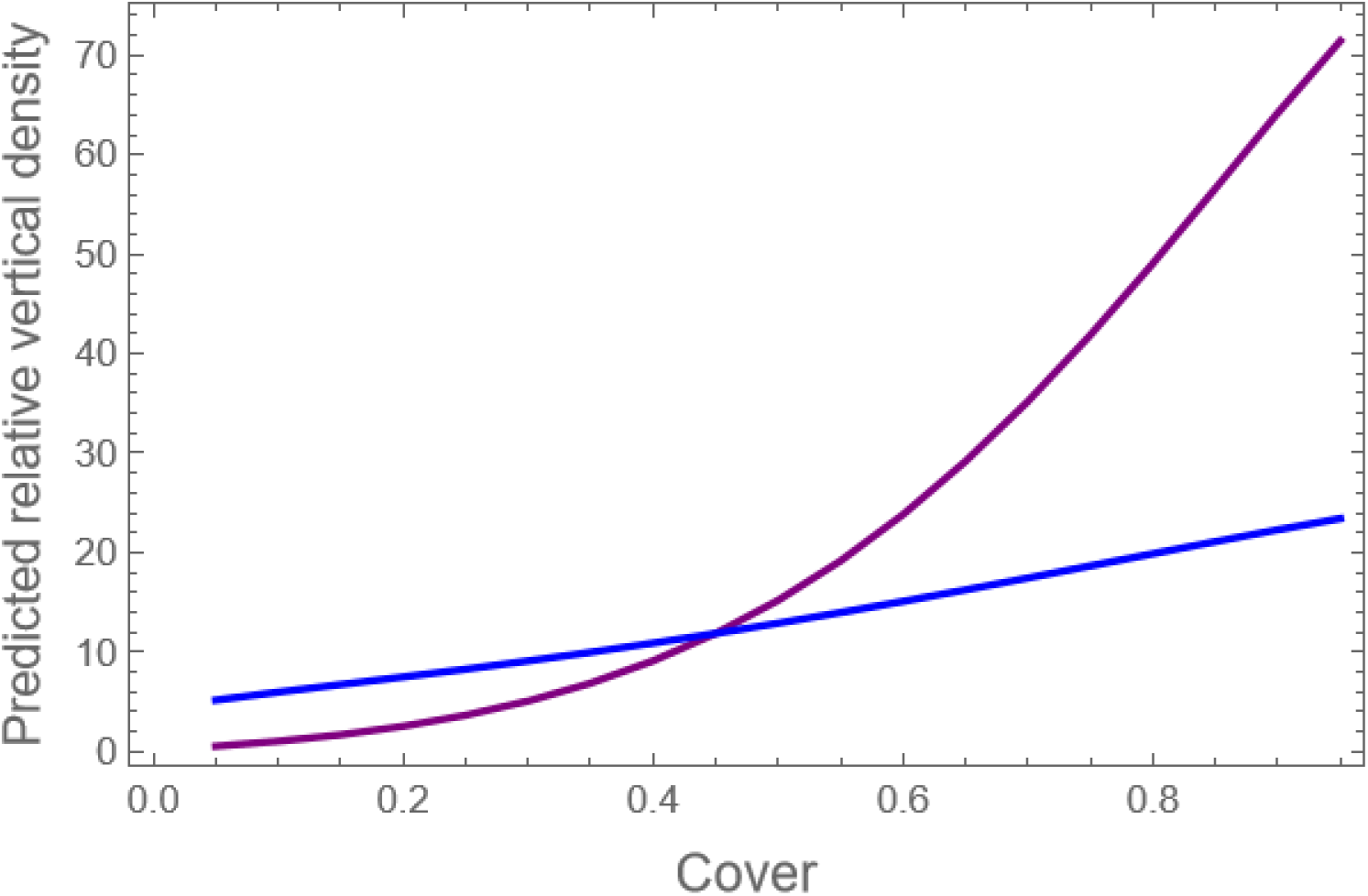
The predicted relative vertical density of *Calluna vulgaris* (purple) and *Avenella flexuosa* (blue) as a function of its cover without added nitrogen. The sum of the cover of the two species is set to one, i.e., only these two species are assumed to be locally present.

For *C. vulgaris* and *A. flexuosa*, there was no significant effect of unmeasured environmental variables (Table 2, ρ_1_, ρ_2_), i.e. the covariance between the early and later growth stages is assumed to be accounted for by the competition model (eq. 1) and not by additional unmeasured variables. However, for all other vascular species there was an indication that unmeasured environmental variables had a significant positive effect on the observed covariance between the early and later growth stages that was not accounted for by the species interaction model (Table 2, ρ_3_).

## Discussion

In the dune heath ecosystem studied, we have presented evidence that both density- and frequency-dependent interspecific competition play significant roles in determining plant growth in the plant community studied. Significant interspecific density-dependent competition was observed in 4 out of the 6 possible cases of three species interactions. We expected that nitrogen addition would alter the species interactions, but this was only demonstrated in the effect of *C. vulgaris* on the growth of all other vascular species, where nitrogen addition increased the competitive effect of *C. vulgaris*. It has been suggested that *C. vulgaris* may retain a competitive advantage with increased deposition of nitrogen because of the vertical structure of its leaves, the mature dense vegetation and the fact that leaves remain on the plant during winter (Aerts et al. 1990; Aerts and Heil 1993). It is possible that the effects of nitrogen addition occur only after several years, after which the level of available nitrogen has reached a new and higher steady-state level. It will be interesting to follow the effect of nitrogen on plant species interactions in the coming years.

*C. vulgaris* and *A. flexuosa* showed positive feedback dynamics on growth when they were relatively dominant at the plot scale, whereas the growth of all other vascular species was favored when they were relatively rare. The positive feedback dynamics of *C. vulgaris* could be explained by the resource monopolizing strategy mediated by ericoid mycorrhiza (Diemont and Heil 1984; Johansson 2001; Smith and Read 2008; Talbot et al. 2008), although the estimated positive feedback was found to increase with added nitrogen for both species, which is somewhat surprising in light of the resource monopolizing hypothesis. On the other hand, there was a beneficial effect of being relatively rare on the growth of all other vascular species in plots when nitrogen was added. This advantage of being rare could be due to the absence of negative interactions with soil pathogens or other natural enemies (e.g. van der Putten et al. 2013). This hypothesis is consistent with the finding of positive covariation between early and late season growth (Rinella et al. 2020). It has been shown that soil inoculation significantly affects heathland plant communities (van der Bij et al. 2017; Wubs et al. 2016) and that the composition of fungal communities depends on the composition of the plant community (Radujković et al. 2020), suggesting that plant–soil biotic interactions are especially important in heathland ecosystems. However, since the aggregated group “other plant species” is made up of many species, the estimated effects will be the net result of different species effects, and the effects of individual species is unknown. It would require a larger experimental set-up to determine the density- and frequency-dependent competition effects of each individual species.

For the two dominant species and the group of remaining species, the results show significant frequency-dependent effects with positive effects of either being relatively dominant or relatively rare.

Implicitly, it may be assumed that biomass at the end of the growing season translates to cover at the start of the next growing season in a species-specific way, so that the observed plant growth patterns may be used to predict community dynamics (Damgaard 2011; Damgaard et al. 2009). The demonstrated positive effects of being relatively dominant for *C. vulgaris* and *A. flexuosa* at the plot scale are corroborated by previous studies, where it was concluded that the more locally dominant of the two species will outcompete the other (Damgaard et al. 2009). The observed advantage of being dominant is also in agreement with divergent long-term successional patterns on dry heath plots, where plots either became dominated by grass or dwarf shrubs (Ransijn et al. 2015).

The observed widespread infestation of the heather beetle in 2020 had adverse effects on *C. vulgaris* cover and growth. Such infestations may play an important role in transforming a dwarf shrub-dominated heath to a grass-dominated heath. However, our results suggest that a single heather beetle outbreak does not have long-term effects on the growth and competitive ability of heather at this dune heath site (Kaae et al. 2024). It appears that *C. vulgaris* is able to recover relatively quickly after infestation events, although recurrent outbreaks could result in a transformation to a grass-dominated heath.

As previously demonstrated (Damgaard and Weiner 2021), there is compelling evidence that the standard use of the Lotka-Volterra model in studies of species interactions in plant communities is inadequate, and that most species interaction studies would be improved by including frequency-dependency. It has also been demonstrated that if the effects of measurement errors are ignored, model predictions may be biased (Carroll et al. 2006; Damgaard 2020; Damgaard and Weiner 2021; Detto et al. 2019). It is therefore also generally advisable to consider the possible effect of measurement errors when modelling plant species interactions.

Traditional vegetation sampling in relatively small plots is not ideal for studying patch dynamic mechanisms, such as positive feedback mechanisms, and cyclical vegetation processes with ageing, which occur at a larger spatial scale. More research on the combined effects of density- and frequency-dependent competition at larger spatial scales is needed. New drone-based sampling methods of plant abundance that operate at a larger spatial scales (e.g. Damgaard 2021) may provide much needed information for the study of plant patch dynamics.

In conclusion, including frequency-as well as density-dependent effects, and unmeasured variables in models of plant community dynamics could result in important improvements in our understanding of plant growth, plant community dynamics and coexistence theory. Modern statistical modelling methods make this easy to do.

## Acknowledgements

We are grateful for funding from Aage V. Jensens Fonde and the Danish Long-term Ecosystem Research Network (LTER-DK).

## Electronic Supplements

The electronic supplements are located at https://osf.io/n6s9w/

Appendix A: Deer densities at Vust heath

Appendix B: Pin-point data (Excel file)

Appendix C: Software code (Mathematica notebook, get free viewer at https://www.wolfram.com/player/)

Fig. S1: Residual plots of vertical density

